# Avian and Human Influenza Viruses Exhibit Distinct Glycoconjugate Receptor Specificities in Human Lung Cells

**DOI:** 10.1101/2022.06.29.498208

**Authors:** Chieh-Yu Liang, Iris Huang, Julianna Han, Senthamizharasi Manivasagam, Jesse Plung, Miranda Strutz, Yolanda Yu, Matheswaran Kandasamy, Francoise A. Gourronc, Aloysius J. Klingelhutz, Biswa Choudhury, Lijun Rong, Jasmine T. Perez, Sriram Neelamegham, Balaji Manicassamy

## Abstract

IAV utilize sialic acid (Sia) containing cell surface glycoconjugates for host cell infection, and IAV strains from different host species show preferences for structurally distinct Sia at the termini of glycoconjugates. Various types of cell surface glycoconjugates (N-glycans, O-glycans, glycolipids) display significant diversity in both structure and carbohydrate composition. To define the types of glycoconjugates that facilitate IAV infection, we utilized the CRISPR/Cas9 technique to truncate different types of glycoconjugates, either individually or in combination, by targeting glycosyltransferases essential to glycan biosynthesis in a human lung epithelial cell line. Our studies show that both human and avian IAV strains do not display strict preferences for a specific type of glycoconjugate. Interestingly, truncation of all three types of glycoconjugates significantly decreased the replication of human IAV strains, yet did not impact the replication of avian IAV strains. Taken together, our studies demonstrate that avian IAV strains utilize a broader repertoire of glycoconjugates for host cell infection as compared to human IAV strains.

**Author Summary:** It is well known that influenza A viruses (IAV) initiate host cell infection by binding to sialic acid, a sugar molecule present at the ends of various sugar chains called glycoconjugates. These glycoconjugates can vary in chain length, structure, and composition. However, it remains unknown if IAV strains preferentially bind to sialic acid on specific glycoconjugates for host cell infection. Here, we utilized CRISPR gene editing to abolish sialic acid on different glycoconjugate types in human lung cells, and evaluated human versus avian IAV infections. Our studies show that both human and avian IAV strains can infect human lung cells by utilizing any of the three major sialic acid-containing glycoconjugate types, specifically N-glycans, O-glycans, and glycolipids. Interestingly, simultaneous elimination of sialic acid on all three glycoconjugate types in human lung cells dramatically decreased human IAV infection, yet had little effect on avian IAV infection. Our studies indicate that avian IAV strains can utilize a wide variety of glycoconjugates for infection, whereas human IAV strains display restrictions in glycoconjugate type usage. These novel studies show distinct differences in host glycoconjugate preferences between human and avian IAV strains.

## Introduction

Host glycans expressed on the cell surface in the form of glycoconjugates (glycoproteins, glycolipids, and glycosaminoglycans) serve as the main entry receptor(s) for a variety of viruses (Kuchipudi et al., 2021; Palese P and Shaw ML, 2007). This is exemplified by influenza A viruses (IAV), which utilize sialic acid (Sia) as the host entry receptor. Sia belongs to a family of 9-carbon sugars (>90 members) predominantly present at the termini of cell surface glycoconjugates in the deuterostome lineage, including vertebrates. As such, IAV can infect a broad range of species, including humans, aquatic birds, domestic birds, swine, and sea mammals, with aquatic birds serving as the reservoir species for almost all subtypes of IAV (Kessler et al., 2021; Webster et al., 1992). In humans and pigs, IAV infections occur in the upper respiratory tract, whereas IAV infections occur in the gastrointestinal tract of avian species. IAV strains from various host species show preferences for distinct modifications on Sia molecules in the context of their linkages and backbone sugar chains (Karakus et al., 2020). Human IAV strains preferentially bind to Sia linked to the penultimate galactose via a α2,6 carbon linkage (*i.e.* Siaα2-6Galβ), whereas avian strains prefer α2,3 linked Sia moieties (Raman et al., 2014; Shi et al., 2014). Differences in IAV host tissue tropism have been attributed to the availability of different Sia types, as Siaα2,6Gal is abundant in the human upper respiratory tract and Siaα2,3Gal is highly expressed in the avian intestinal tract (de Graaf and Fouchier, 2014). As both types of sialoglycans are expressed in the respiratory tract of swine, they are able to support the replication of both avian and human IAV strains (de Graaf and Fouchier, 2014). In the past 100 years, IAV strains from zoonotic reservoirs have crossed the species barrier and caused four pandemics in humans. These pandemic strains demonstrate the unique ability to bind to Sia receptors present in both human and avian hosts (Shi *et al*., 2014; Stevens et al., 2006). Thus, the ability to bind Siaα2,6Gal versus Siaα2,3Gal is considered a critical factor in determining IAV host range (Shi *et al*., 2014).

Sia containing glycoconjugates are attached to proteins through asparagine (N-glycan) or serine/threonine residues (O-glycan), or to glycosphingolipids (GSL). IAV entry is initiated by binding of viral hemagglutinin (HA) to cell surface sialoglycans, which triggers intracellular signaling cascades, such as receptor tyrosine kinases (RTK), that facilitate virion uptake and fusion (Eierhoff et al., 2010). Prior studies suggest that HA can engage Sia modifications present on several cell surface proteins, such as epidermal growth factor receptor (EGFR), calcium-dependent voltage channel (Ca_v_1.2i), natural killer cell receptors (NKP44/46), and nucleolin, to facilitate virion uptake (Karakus *et al*., 2020). Moreover, in the absence of Sia, some C-type lectins predominantly expressed on antigen presenting cells, such as DC-SIGN, L-SIGN, mannose receptor, etc., can also facilitate IAV uptake by binding to glycan moieties on viral glycoproteins (Karakus *et al*., 2020). The glycoconjugate structural features necessary for HA binding have been identified using chemically defined glycan arrays (Consortium for Functional Glycomics - CFG) and shotgun lung tissue glycan arrays (Byrd-Leotis et al., 2019b; Byrd-Leotis et al., 2014; Connor et al., 1994; Jia et al., 2020; Rogers and Paulson, 1983; Stevens *et al*., 2006). These studies suggest that human IAV strains preferentially bound to long branched sialoglycans with poly-lactosamine (polyLN) repeats that had an ‘umbrella-like’ topology, whereas avian IAV strains preferentially bound to short sialoglycans with a single lactosamine that had a ‘cone-like’ topology, indicating that glycan topology can also determine host range (Chandrasekaran et al., 2008). In agreement, circulating human H3N2 viruses have evolved to utilize extended branched polyLN glycans (N-glycans), while the parental pandemic H3N2 strain preferentially bound to short sialyl-LN glycans (Broszeit et al., 2021; Byrd-Leotis et al., 2019a; Peng et al., 2017). In array slides, glycans are immobilized in non-natural configurations at a high density with uniformity and hence, the conclusions can be biased on the repertoire of glycans presented (Raman *et al*., 2014; Shi *et al*., 2014).

Two prior studies using Chinese hamster ovary (CHO) cell lines lacking N-glycans (due to mutations in an essential biosynthesis gene Mgat1) arrived at opposite conclusions - N-glycans were absolutely required for IAV infection (Chu and Whittaker, 2004), and N-glycans were not an absolute requirement (de Vries et al., 2012). The results from the latter study are consistent with binding studies performed using recombinant HA and a panel of human embryonic kidney (HEK) 293 CRISPR knock out cells (Narimatsu et al., 2019). In addition, studies in HEK293 CRISPR KO cells also reconfirmed the Siaα2,3 versus Siaα2,6 binding preferences for avian and human HAs, respectively (Narimatsu *et al*., 2019). However, these studies were limited to the assessment of HA binding; IAV infection and replication were not evaluated. Importantly, as HEK 293 cells likely do not mimic the glycan repertoire of human lung epithelial cells, our understanding of the types of glycoconjugates utilized by human and avian IAV strains for infection of human lung cells remains incomplete.

In this study, we utilized a human lung epithelial cell line (A549) to assess the preferences for different glycoconjugate types by avian versus human IAV strains in the context of infection. To this end, a comprehensive panel of CRISPR gene edited A549 cells were generated that contained truncated N-glycans [N]^-^, O-glycans [O]^-^, or glycosphingolipids [G]^-^, either individually or in combination, by disrupting the expression of glycosyltransferases *MGAT1*, *C1GALT*, or *UGCG*, respectively. Surprisingly, truncation of individual glycans ([N]^-^, [O]^-^, [G]^-^) in A549 cells had no effect on the replication of multiple IAV strains; concurrent truncation of two glycan types ([NO]^-^, [NG]^-^) in A549 cells showed a modest decrease in H1N1 infection, yet no defect in avian H5N1 infection. Thus, beyond the known Siaα2,3 versus Siaα2,6 differences, the glycoconjugate types also determine the functional diversity of IAV strains. Importantly, concurrent truncation of all 3 glycan types ([NOG]^-^) in A549 cells resulted in a 1-3 log decrease in viral titers for human H1N1 and H3N2 viruses, yet showed little to no change in titers for several avian IAV strains (H5N1, H7N7, H7N9, H7N9 etc.). The robust replication of avian IAV strains observed in A549 ([NOG]^-^) cells was dependent on Sia receptors, suggesting that glycoconjugates may also contribute to infection. Taken together, our study is the first to demonstrate the functional relevance of different glycoconjugate types for avian versus human IAV strains. Importantly, our studies show that avian IAV strains utilize structurally diverse glycoconjugates for host cell infection as compared to human IAV strains.

## Results

### Generation of A549 cells lacking Sia on N- or O-glycans

We ablated terminal Sia modifications on specific glycoconjugate types in human lung epithelial A549 cells by targeting glycosyltransferases that are essential for the elongation of cell-surface N- or O-linked glycans (Figure 1A) (Stolfa et al., 2016). Here, using the CRISPR/Cas9 technology, we targeted the glycosyltransferases *MGAT1* or *C1GALT1* to generate A549 cells with truncated N-glycans ([N]^-^) or O-glycans ([O]^-^), respectively (Figure 1B and Table S1). Loss of individual glycosyltransferases was confirmed by sanger sequencing of the sgRNA target site, western blot analysis, and lectin staining with *Sambucus Nigra* lectin (SNA; specific for α2,6-linked Sia), *Vicia Villosa* lectin (VVL, specific for terminal GalNAc), and Cholera toxin B subunit (CTB, specific for GM1 gangliosides) (Figure S1 and Table S2). As previously described, we observed decreased SNA binding in *MGAT1* KO cells ([N]^-^) as compared to wild type (WT) A549 cells, demonstrating that truncation of N-glycan structures resulted in the loss of N-glycans with α2,6-linked Sia moieties (Figure S1C) (Stolfa *et al*., 2016). As expected, we observed similar levels of VVL and CTB binding in *MGAT1* KO cells ([N]^-^) as compared to WT cells, indicating that the expression of O- and GSL-glycans was not significantly altered upon loss of MGAT1. In *C1GALT1* KO cells ([O]^-^), we observed increased VVL binding due to higher levels of unmodified GalNAc on truncated O-glycans (Figure S1C). As expected, *C1GALT1* KO cells ([O]^-^) showed similar levels of SNA and CTB binding as compared to WT cells, indicating that the expression of N- and GSL-glycans remained unaltered. Taken together, we successfully generated A549 cells lacking Sia specifically on N- or O-glycans.

**Figure 1.**
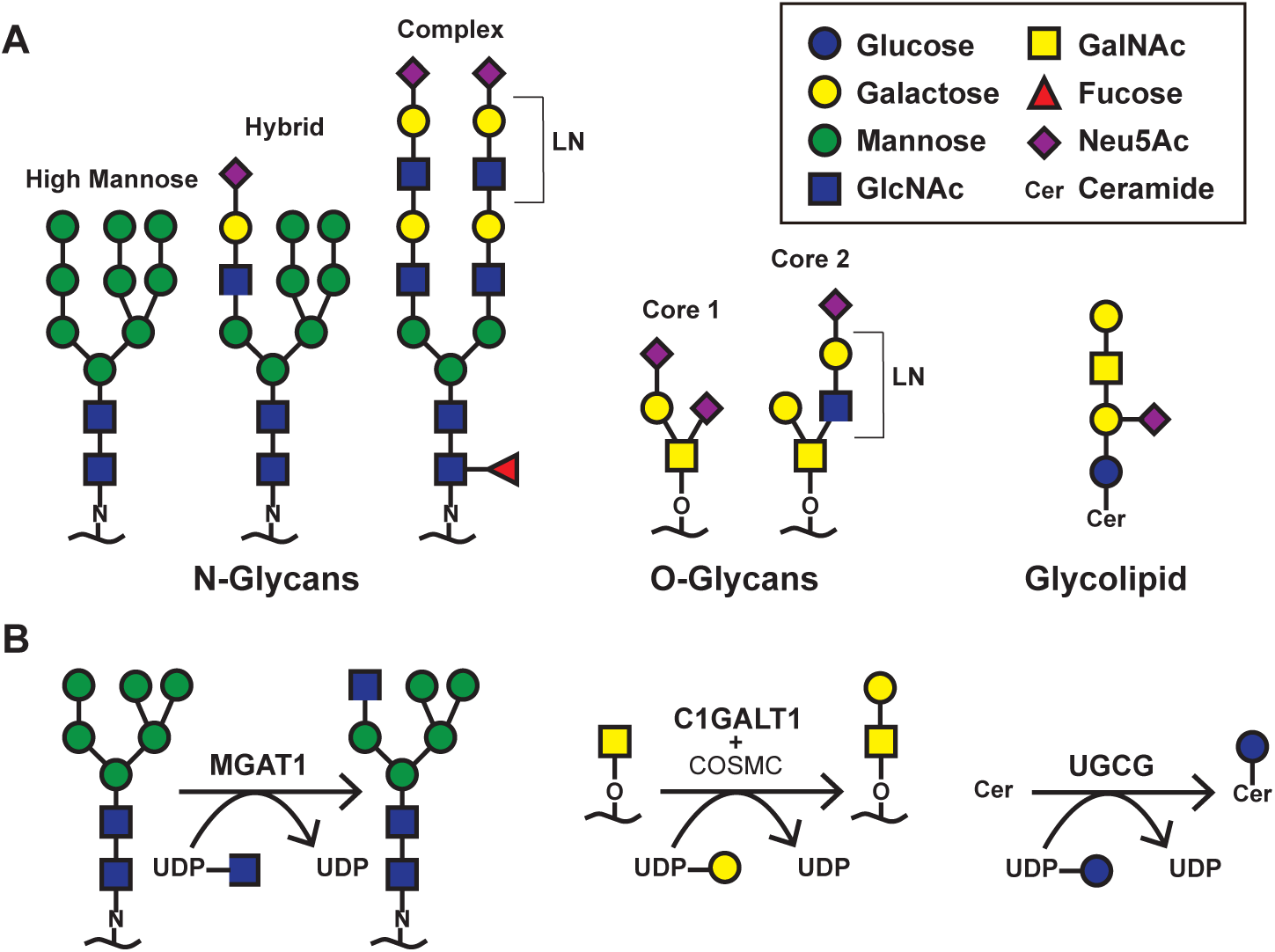
Representative structures of Sia-containing glycoconjugates and the key glycosyltransferases essential for glycoconjugate biosynthesis. (A) Schematic representation of different glycans terminating with Sia. For N-glycans: high-mannose, hybrid, and complex structures are shown; for O-glycans: Core 1 and Core 2 structures are shown; ganglioside GM1 type glycolipid is shown. (B) Key glycosyltransferases essential for biosynthesis of individual glycan types. MGAT1 is necessary for the formation of hybrid and complex N-glycans. C1GALT1 is essential for the synthesis of Core 1 and Core 2 O-glycans. UGCG is essential for the first step of glycolipid biosynthesis.

### Sia-containing N-glycans or O-glycans are not essential for IAV replication

To determine if removal of terminal Sia on either N- or O-glycans impaired IAV infection, we performed single-cycle infection assays in [N]^-^ and [O]^-^ KO cells with H1N1 (H1N1-GFP) and H5N1 (H5N1-GFP) viruses at a high multiplicity of infection (MOI) (MOI=3), and assessed GFP expression at 16 hours post infection (hpi) by flow cytometry. We observed modest differences in the percentage of GFP positive cells between infected WT cells and [N]^-^ or [O]^-^ KO cells, indicating that truncation of N- or O-glycans individually did not grossly affect single-cycle IAV infection (Figure 2A). Next, we performed multi-cycle replication assays in [N]^-^ and [O]^-^ KO cells with four different IAV strains at a low MOI (MOI=0.01- 0.001) and observed robust replication for all four IAV strains in both [N]^-^ and [O]^-^ KO cells at levels similar to WT A549 cells (Figure 2B). Similarly, the replication of vesicular stomatitis virus (VSV), which enters host cells through interactions with the low-density lipoprotein-receptor (LDLR) and thus would be unimpaired by truncation of sialoglycans, also remained unaffected in these KO cells (Finkelshtein et al., 2013). Taken together, these results demonstrate that loss of Sia on N- or O-glycans did not affect IAV infection or replication, indicating that they are not solely essential for IAV entry.

**Figure 2.**
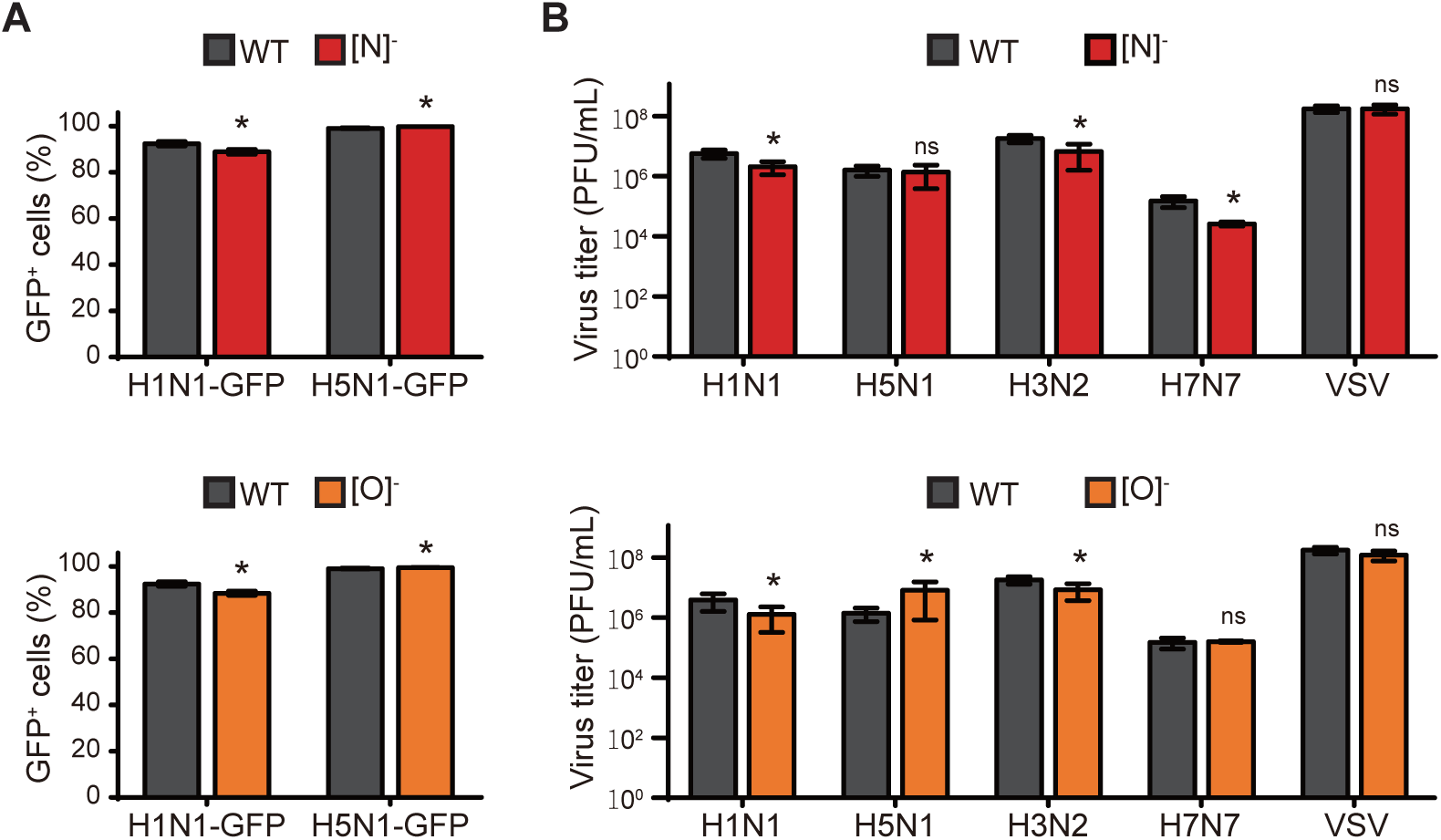
Truncation of N- or O-glycans individually does not affect IAV infection and replication. (A) Single-cycle infection assays with H1N1-GFP and H5N1-GFP in individual glycan KO cells. WT and KO cells seeded in 6-well dishes were infected with H1N1-GFP or H5N1-GFP at a high MOI (MOI=3) without TPCK-treated trypsin and at 16hpi, the percentage (%) of GFP expressing cells was determined by flow cytometric analysis. Top: Comparison of infection levels between WT and [N]^-^ KO cells; bottom: comparison of infection levels between WT and [O]^-^ KO cells. Data are represented as mean percentage of GFP+ cells from triplicate samples ± SD. (B) Multi-cycle replication assays with various IAV strains and VSV in individual glycan KO cells. WT and KO cells seeded in 6-well dishes were infected with various IAV strains or VSV at a low MOI (MOI=0.001-0.01) in the presence of TPCK-treated trypsin and at 48hpi, viral titers in the supernatants were determined by plaque assay in MDCK cells. Top: comparison of viral replication in WT and [N]^-^ KO cells; bottom: comparison of viral replication in WT and [O]^-^ KO cells. Data are represented as mean titer of triplicate samples ± SD (PFU/mL). * denotes p-value ≤0.05. ns is non-significant. Data are representative of at least three independent experiments.

### Sia-containing O-glycans or glycosphingolipids can individually support robust IAV replication

Next, we generated A549 double knockout (DKO) cells with truncations in both N- and O-glycans ([NO]^-^ DKO; expressing only glycosphingolipids (GSL)) as well as DKO cells with truncations in both N-glycans and GSL ([NG]^-^ DKO; expressing only O-glycans), and confirmed truncation of the intended glycans as described above (Figure S2A-B and Table S2). Along with the previously described lectins (SNA, VVL, and CTB), we also assessed binding of *Galanthus Nivalis* lectin (GNL), which shows affinity for high mannose containing N-glycans. We observed increased GNL binding in both [NO]^-^ and [NG]^-^ DKO cells as compared to WT cells, as loss of MGAT1 increases the levels of high mannose containing N-glycans (Figure S2B) (Stolfa *et al*., 2016). In addition, we confirmed the lack of SNA binding in both [NO]^-^ and [NG]^-^ DKO cells. The disruption of O-glycans or GSL structures in [NO]^-^ and [NG]^-^ DKO cells was verified by increased VVL binding and decreased CTB binding, respectively.

To determine if concurrent truncation of two glycan types impairs IAV infection, we performed single-cycle infections in [NO]^-^ and [NG]^-^ DKO cells with H1N1-GFP or H5N1- GFP. We observed a 30-40% decrease in H1N1-GFP infection for both [NO]^-^ and [NG]^-^ DKO cells as compared to WT cells (Figure 3A); in contrast, H5N1-GFP infection remained high in both DKO cell types at levels comparable to WT cells. Next, we performed multi-cycle replication assays in [NO]^-^ and [NG]^-^ DKO cells with H1N1 or H5N1 viruses (Figure 3B and Figure S2B). Surprisingly, replication of both IAV strains remained high in [NO]^-^ and [NG]^-^ DKO cells, with peak viral titers reaching up to ∼10^7^ PFU/mL at 48hpi. These results demonstrate that Sia on a single major glycoconjugate type (O-glycans or GSL) is sufficient to support robust replication of H1N1 and H5N1, albeit with reduced H1N1-GFP infection.

**Figure 3.**
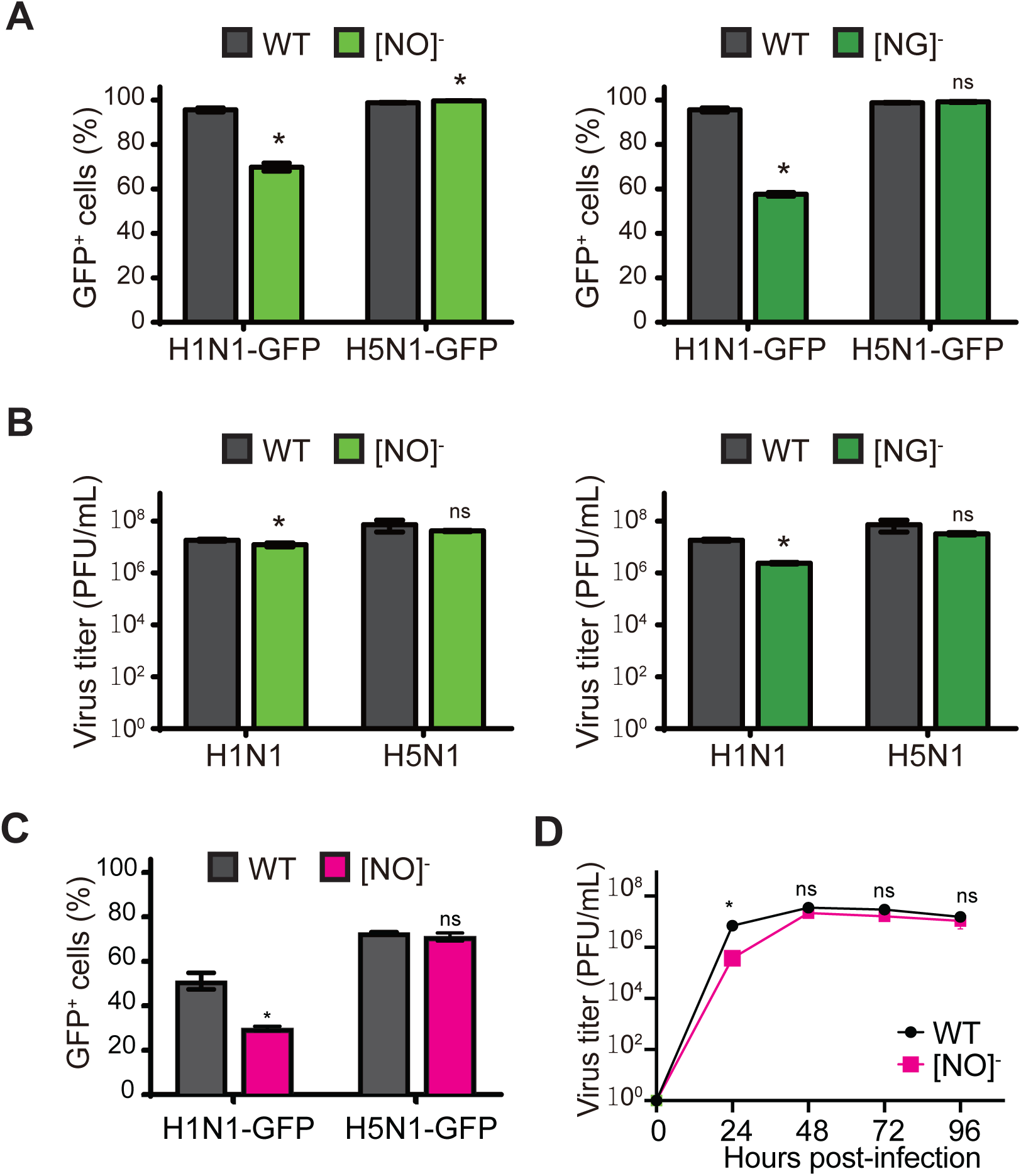
Concurrent truncation of two glycan types decreases H1N1 but not H5N1 infection. (A) Single-cycle infection assays with H1N1-GFP and H5N1-GFP in [NO]^-^ and [NG]^-^ DKO cells. WT and DKO cells seeded in 6-well dishes were infected with H1N1-GFP or H5N1-GFP at a high MOI (MOI=3) without TPCK-treated trypsin and at 16hpi, the % of GFP expressing cells was determined by flow cytometry analysis. Left: comparison of infection levels between WT and [NO]^-^ DKO cells; right: comparison of infection levels between WT and [NG]^-^ DKO cells. Data are presented as mean percentage of GFP+ cells from triplicate samples ± SD. (B) Multi-cycle replication assays with H1N1 and H5N1 viruses in [NO]^-^ and [NG]^-^ DKO cells. WT and DKO cells seeded in 6-well dishes were infected at a low MOI with H1N1 (MOI=0.01) or H5N1 (MOI=0.001) in the presence of TPCK-treated trypsin and at 48hpi, viral titers in the supernatants were determined by plaque assay in MDCK cells. Left: comparison of viral replication in WT and [NO]^-^ DKO cells; right: comparison of viral replication in WT and [NG]^-^ DKO cells. (C) Single-cycle infection assays with H1N1-GFP and H5N1-GFP in Nuli-1 [NO]^-^ DKO cells. Nuli-1 WT and DKO cells seeded in 6-well dishes were infected with H1N1-GFP or H5N1-GFP at a high MOI (MOI=3) without TPCK-treated trypsin and at 16hpi, the % of GFP positive cells was determined by flow cytometric analysis. Data are represented as mean percentage of GFP+ cells from triplicate samples ± SD. (D) Multi-cycle replication assays with H5N1 virus in Nuli-1 [NO]^-^ DKO cells. Nuli-1 WT and DKO cells seeded in 6-well dishes were infected at a low MOI with H5N1 (MOI=0.001) in the presence of TPCK-treated trypsin and viral titers in the supernatants at different hpi were determined by plaque assay in MDCK cells. Data are represented as mean titer of triplicate samples ± SD (PFU/mL). * denotes p-value ≤0.05. ns is non-significant. Data are representative of at least three independent experiments.

### Validation of IAV glycoconjugate requirements in Nuli-1 cells

To validate our findings in another human lung cell line, we investigated the importance of different glycoconjugate types for IAV replication in Nuli-1 cells. We generated Nuli-1 DKO cells with concurrent truncation of both N- and O-glycans by targeting the *MGAT1* and *C1GALT1* genes via CRISPR editing and confirmed loss of the intended glycans as described above (Nuli-1 [NO]^-^ DKO cells; Fig S2D). Similar to our findings in A549 [NO]^-^ DKO cells, we observed reduced H1N1-GFP infection in Nuli-1 [NO]^-^ DKO cells as compared to control Nuli-1 cells (Figure 3C). In contrast, H5N1-GFP infection in Nuli-1 [NO]^-^ DKO cells remained high, with levels similar to control Nuli-1 cells. In addition, in multi-cycle replication assays, H5N1 replication in Nuli-1 [NO]^-^ DKO cells was comparable to control Nuli-1 cells (Figure 3D). Unlike H5N1, H1N1 showed poor replication in control Nuli-1 cells and hence, we did not perform multicycle replication assays with H1N1. Taken together, our studies in Nuli-1 [NO]^-^ DKO cells confirmed our findings in A549 cells that Sia on GSL is sufficient for robust H5N1 replication.

### Generation and characterization of A549 cells lacking Sia on three glycoconjugate types

Next, we generated A549 triple KO (TKO) cells with truncations in N-glycans, O-glycans, and GSL ([NOG]^-^ TKO) and confirmed loss of the intended glycan types as described above (Figure S3A-B and Table S2). As anticipated, [NOG]^-^ TKO cells showed increased GNL binding, decreased SNA binding, increased VVL binding, and decreased CTB binding (Figure S3B). To define the structural features of N- and O-glycans expressed in [NOG]^-^ TKO cells, we performed high pH anion exchange chromatography (HPAEC) followed by fluorescence detection (HPAEC-FL) as well as matrix-assisted laser desorption/ionization time-of-flight/time-of-flight (MALDI-TOF/TOF) mass spectrometry (Figures S3C and 4). In WT A549 cells, we detected bi- and tri-antennary N-glycan structures terminating in high-mannose or Sia by both techniques. In contrast, we only detected N-glycan structures terminating in high mannose residues in [NOG]^-^ TKO cells by both techniques. In our O-glycan profiling of WT A549 cells, we observed Core 1 and Core 2 structures with or without terminal sialic acid modifications (Figure 4B); in [NOG]^-^ TKO cells, we did not observe distinct O-glycan structures. As GSL are less abundant in A549 cells, we were unable to perform mass spectrometry analysis to confirm the loss of GSL in [NOG]^-^ TKO cells. Both CTB binding assay and sanger sequencing of the sgRNA target region confirmed the loss of UGCG in [NOG]^-^ TKO cells (FigureS3B and Table S2). Together, these results confirmed truncation of these three glycan types in [NOG]^-^ TKO cells.

**Figure 4.**
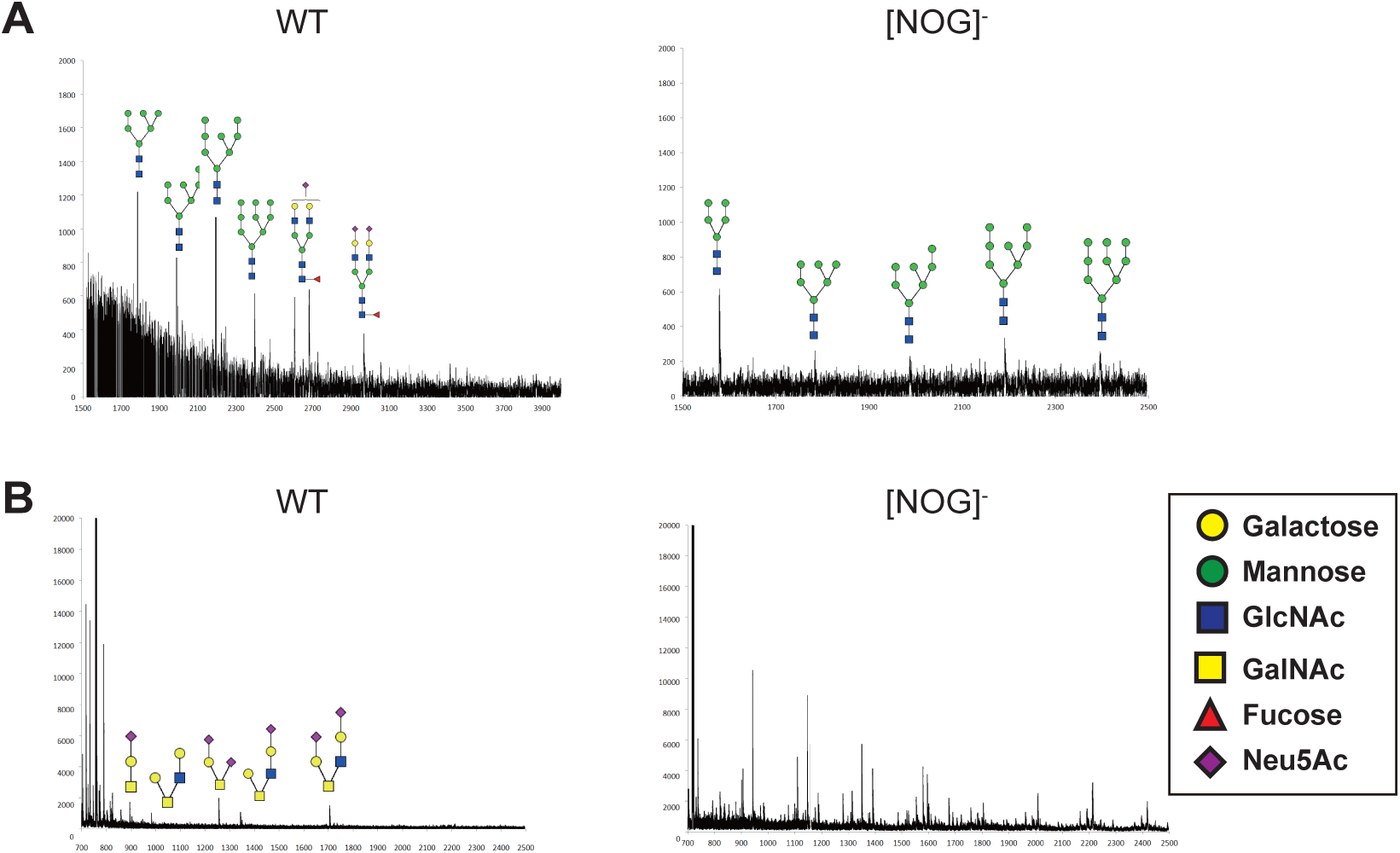
MALDI TOF/TOF analysis for N- and O-glycan structures in A549 and [NOG]^-^ TKO cells. N-glycans and O-glycans were extracted from uninfected A549 WT and [NOG]^-^ TKO cells and subjected to MALDI TOF/TOF analysis. (A) Comparison of N-glycans expressed in WT and [NOG]^-^ TKO cells. (B) Comparison of O-glycans expressed in WT and [NOG]^-^ TKO cells.

### H1N1 requires Sia on one of the three major glycoconjugates for robust replication

Next, we compared single-cycle infections of H1N1-GFP and H5N1-GFP in [NOG]^-^ TKO and WT cells. H1N1-GFP infection was drastically reduced by >95% in [NOG]^-^ TKO cells as compared to WT cells (Figure 5A). Surprisingly, we only observed an ∼20% reduction in H5N1-GFP infection in [NOG]^-^ TKO cells as compared to WT cells. To confirm that the observed reduction in IAV infection was due to decreased virus binding, we performed cell surface binding assays with purified HA proteins and IAV. In HA binding assays, both H1 and H5 HA proteins showed reduced binding in [NOG]^-^ TKO cells as compared to WT cells (Figure 5B). Interestingly, H5N1 virions showed higher cell surface binding as compared to H1N1 virions in [NOG]^-^ TKO cells (Figure 5C); however, the levels of binding for both H1N1 and H5N1 virions were lower in [NOG]^-^ TKO cells as compared to WT cells. To confirm these results, we performed single-cycle high MOI kinetics assays and observed reduced H1N1 virion production over time from [NOG]^-^ TKO cells as compared to WT cells (Figure 5D). Next, we performed multi-cycle replication assays and observed a 2-3 log decrease in viral titers over time for H1N1 in [NOG]^-^ TKO cells as compared to WT cells (Figure 5E). In contrast, H5N1 virus showed robust replication in [NOG]^-^ TKO cells, with only a modest decrease in viral titers in [NOG]^-^ TKO cells as compared to WT cells. Together, these data indicate that truncation of three glycan types decreased the susceptibility of [NOG]^-^ cells to H1N1 infection, yet did not dramatically affect H5N1 infection.

**Figure 5.**
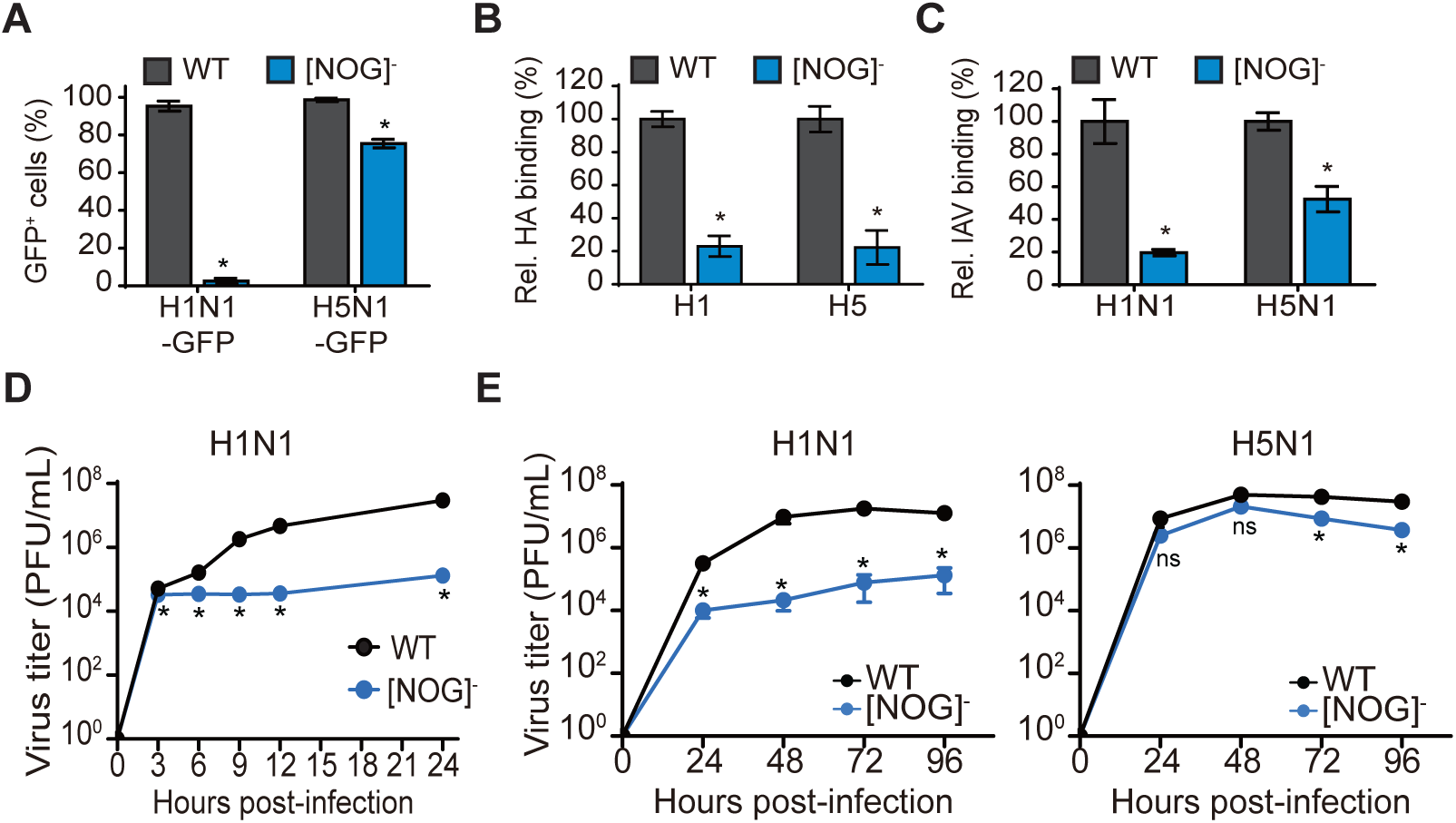
H5N1 but not H1N1 shows robust infection and replication in [NOG]^-^ TKO cells. (A) Single-cycle infection assays with H1N1-GFP and H5N1-GFP in [NOG]^-^ TKO cells. WT and TKO cells seeded in 6-well dishes were infected with H1N1-GFP or H5N1-GFP at a high MOI (MOI=3) without TPCK-treated trypsin and at 16hpi, the % of GFP positive cells was determined by flow cytometric analysis. Data are represented as mean % of GFP+ cells from triplicate samples ± SD. (B) Comparison of cell surface binding of purified HA in [NOG]^-^ TKO cells. WT or [NOG]^-^ TKO cells were incubated with purified H1 or H5 subtype HA on ice and HA binding was measured by flow cytometry. (C) Comparison of cell surface binding of H1N1 and H5N1 virions in [NOG]^-^ TKO cells. WT or [NOG]^-^ TKO cells were incubated with H1N1 or H5N1 virus (MOI=100) on ice and virion binding was measured by flow cytometry. For B and C, data are represented as mean relative binding from triplicate samples ± SD. (D) Single-cycle replication assays with H1N1 in [NOG]^-^ TKO cells. WT and DKO cells seeded in 6-well dishes were infected at an MOI=3 without TPCK-treated trypsin and at various times post-infection, supernatants were collected and viral titers were determined after the addition of TPCK-treated trypsin 1hr prior to plaque assay. (E) Multi-cycle replication assays with H1N1 and H5N1 in [NOG]^-^ TKO cells. WT and DKO cells seeded in 6-well dishes were infected at a low MOI with H1N1 (MOI=0.01) or H5N1 (MOI=0.001) in the presence of TPCK-treated trypsin and viral titers in the supernatants at different hpi were determined by plaque assay in MDCK cells. For D and E, data are represented as mean titer of triplicate samples ± SD (PFU/mL). * denotes p-value ≤0.05. ns is non-significant. Data are representative of at least three independent experiments.

### Various avian IAV strains show versatility in Sia receptor usage

Next, we expanded the panel of influenza viruses and tested the replication of a variety of influenza A and B viruses in [NOG]^-^ TKO cells. We observed a 1-3 log decrease in the replication of human and swine H1N1 and H3N2 subtypes as well as influenza B viruses, with the exception of the 1968 pandemic H3N2 virus that originated from an avian host (Figure 6A-B). The 1968 H3N2 strain showed robust replication in [NOG]^-^ TKO cells at levels comparable to WT cells (Figure S4A). Interestingly, avian IAV replication of H4, H5, H7, and H9 subtypes remained high in [NOG]^-^ TKO cells (Figures 6C-D and S4B). Taken together, our findings show that human and swine IAV strains require Sia on one of the three major glycoconjugates for viral entry, whereas avian IAV strains utilize an expanded repertoire of glycoconjugates.

**Figure 6.**
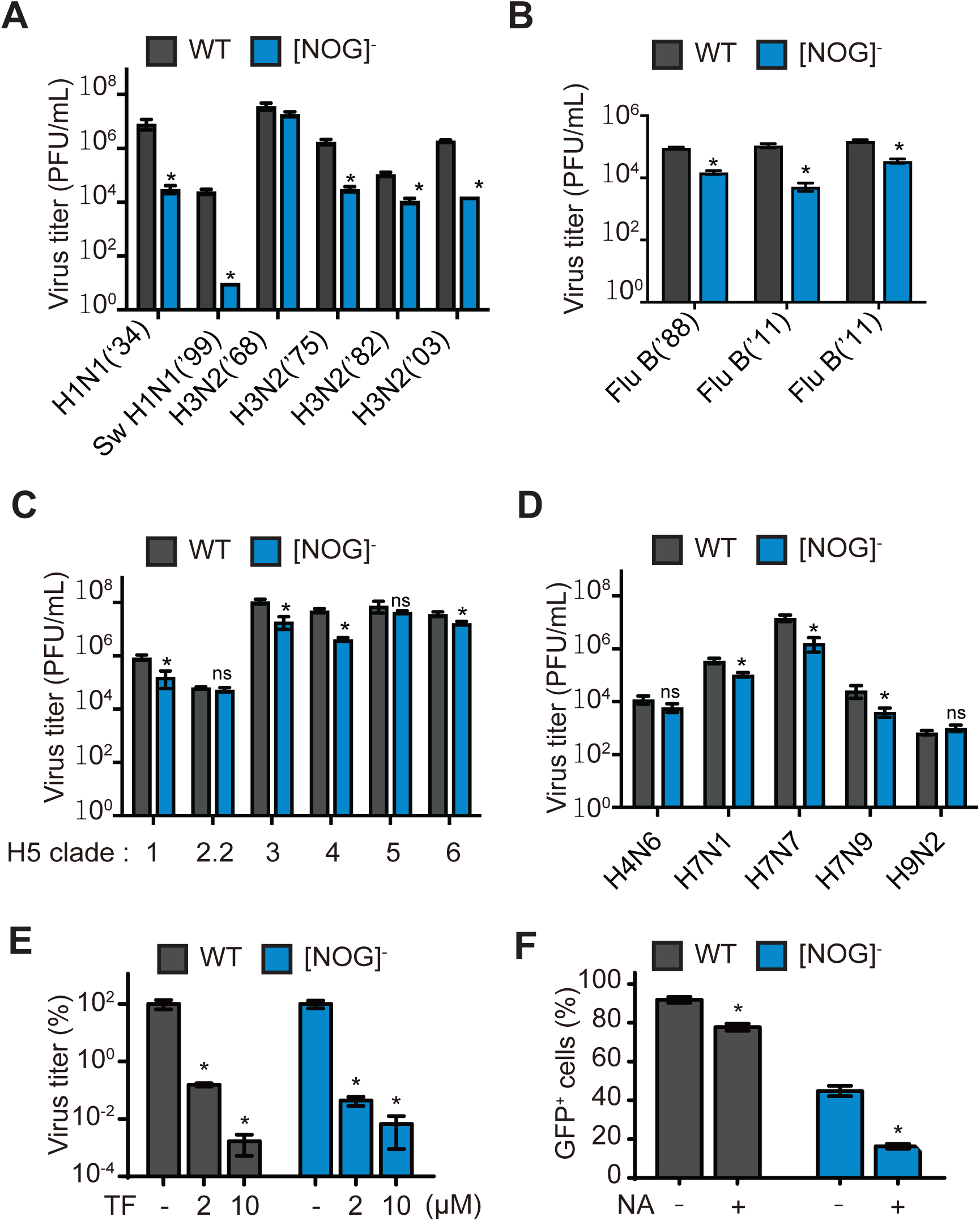
Avian but not human IAV strains show robust replication in [NOG]^-^ TKO cells in a Sia dependent manner. (A-D) Multi-cycle replication assays with various IAV and influenza B strains in [NOG]^-^ TKO and WT cells. (A) Replication of human and swine IAV strains. (B) Replication of human influenza B virus strains. (C) Replication of avian H5N1 viruses from different representative clades. (D) Replication of other avian IAV strains. Data are represented as mean titer of triplicate samples ± SD (PFU/mL). (E) Treatment with Oseltamivir carboxylate limits H5N1 replication in [NOG]^-^ TKO cells. WT and DKO cells seeded in 6-well dishes were infected at a low MOI with H5N1 (MOI=0.001) and incubated with the indicated concentrations of Oseltamivir carboxylate (‘TF’) in the presence of TPCK-treated trypsin. At 48hpi, viral titers in the supernatants were determined by plaque assay in MDCK cells. Data are represented as a percentage mean titer of triplicate samples relative to untreated cells ± SD. (F) Sialidase pretreatment decreases H5N1 infection in [NOG]^-^ TKO cells. WT or [NOG]^-^ TKO cells seeded in 6-well plates were pretreated with 500 mU/mL sialidase from *Clostridium perfringens* (Sigma) for 3 hours at 37°C before infection with H5N1-GFP (MOI=3). At 16hpi, the % of GFP positive cells was determined by flow cytometric analysis. Data are represented as mean percentage of GFP+ cells from triplicate samples ± SD. * denotes p-value ≤0.05. ns is non-significant. Data are representative of at least two independent experiments.

### H5N1 IAV replication in [NOG]^-^ TKO cells is dependent on residual Sia

The ability of several avian IAV strains to replicate in [NOG]- TKO cells led us to consider that there may be residual Sia moieties in [NOG]^-^ TKO cells. As loss of C1GALT1 results in truncation of Core1 and Core 2 O-glycans yet does not impact biosynthesis of sialyl Tn antigen (STn), we assessed STn levels with a specific antibody. We observed increased STn expression in [NOG]- TKO cells as compared to WT A549 cells, suggesting that H5N1 viruses may utilize STn as entry receptors (Figure S5A-B). Next, to demonstrate that avian H5N1 replication occurs in a Sia-dependent manner in [NOG]^-^ TKO cells, we performed multi-cycle replication assays in a presence of viral neuraminidase inhibitor (Oseltamivir carboxylate). In this way, we can determine if IAV virions produced from [NOG]^-^ TKO cells require neuraminidase for release. We observed a >4 log decrease in H5N1 titers upon treatment with Oseltamivir for both [NOG]^-^ TKO cells and WT cells, indicating that viral neuraminidase activity is essential for H5N1 replication in [NOG]^-^ TKO cells (Figure 6E). In addition, pretreatment of [NOG]^-^ TKO cells with *C. Perfringens* sialidase for 2hrs dramatically reduced single-cycle H5N1 infection in [NOG]^-^ TKO cells, validating the Sia-dependent replication of H5N1 in these cells (Figure 6F). Together, our studies demonstrate that avian H5N1 can utilize diverse Sia moieties including residual Sia not present on the three major glycan types.

## Discussion

In this study, we assessed the importance of various cell surface Sia-containing glycoconjugates to human versus avian IAV infection. Using the CRISPR/Cas9 gene editing technique, we truncated the three major types of glycoconjugates (N-glycans, O-glycans, and GSL), either individually or concurrently, in a human lung epithelial cell line (A549) and evaluated IAV replication. Our studies demonstrated that both human and avian IAV strains did not show strict preferences for any of the three types of glycoconjugates for host cell infection. Interestingly, our studies in [NOG]^-^ TKO cells showed that concurrent truncation of the three major glycoconjugates significantly reduced human H1N1 and H3N2 replication, indicating that human IAV strains require the presence of Sia on one of the three major glycoconjugates. In contrast, several avian IAV strains demonstrated robust replication in [NOG]^-^ TKO cells, suggesting that avian IAV strains utilize an expanded repertoire of glycoconjugates. Taken together, our studies reveal that human and swine IAV strains differ starkly from avian IAV strains in glycoconjugate receptor requirements in human lung epithelial cell lines.

It has been well-established that avian and human IAV strains differ in Siaα2,3Gal versus Siaα2,6Gal receptor preferences (Raman *et al*., 2014; Shi *et al*., 2014). However, it remains unknown if a specific type of glycoconjugate serves as the primary receptor or if multiple glycoconjugates are capable of facilitating IAV entry. Much of our understanding of the utilization of glycoconjugate types by various IAV strains has been inferred from *in vitro* virion or purified HA binding assays on glycan array slides (Byrd-Leotis *et al*., 2019b; Byrd-Leotis *et al*., 2014; Connor *et al*., 1994; Jia *et al*., 2020; Rogers and Paulson, 1983; Stevens *et al*., 2006) (Broszeit *et al*., 2021; Byrd-Leotis *et al*., 2019a; Peng *et al*., 2017). Studies with CFG arrays and shotgun lung tissue glycan arrays indicated that human adapted IAV strains preferred longer branched polyLN glycans (N-glycans) for attachment. Interestingly, some of the parental pandemic IAV strains originating from avian hosts bound to short sialyl-LN glycans, suggesting that avian IAV strains may adapt to utilize polyLN in humans. Our mass spectrometry analysis of WT A549 cells showed the presence of short branched N-glycans with one or two LN repeats and single LN containing O-glycans (Figure 4); however, longer branched polyLN were not detected in A549 cells. In congruence with glycan array studies, we observed reduced H1N1-GFP infection in both [NO]^-^ and [NG]^-^ DKO cells, suggesting that human IAV strains may prefer Sia linked to extended glycans with LN repeats (N-glycans) for efficient host cell attachment (Figure 3A). Surprisingly, in the multi-cycle replication studies with H1N1 in [NO]^-^ and [NG]^-^ DKO cells, we observed robust virus replication despite the lack of branched LN repeat containing glycans, indicating that branched LN structures are not absolutely necessary for human IAV infection in A549 cells. In contrast, avian H5N1-GFP infection and H5N1 replication remained grossly unaffected in both [NO]^-^ and [NG]^-^ DKO cells, indicating that shorter glycoconjugates can support robust avian IAV infection. Some of the discrepancies observed between our studies and the aforementioned glycan array studies may be in part due to the manner in which sialoglycans are presented on the host cell surface versus on an array slide (Chan et al., 2013; Raman *et al*., 2014; Shi *et al*., 2014; Walther et al., 2013). In array slides, glycans are presented at a high density with uniformity; as such, the printed glycan arrays may allow for efficient engagement of multiple HA trimers from the same virion. In contrast, HA receptor engagement on the host cell surface likely occurs in a progressive manner through lateral movement of the virion on the cell surface, which may be necessary for activation of intracellular signaling pathways through receptor clustering (Karakus *et al*., 2020). It is possible that in tissue culture, long polyLN containing sialoglycans are not required for IAV infection, yet are necessary for efficient virion binding in glycan arrays. Thus, our CRISPR glycoengineered cells serve as a complementary model system to investigate the importance of host glycans for IAV replication, as the glycans are presented in the context of a host cell.

Similar to the aforementioned glycan array studies, a recent study assessed viral HA binding to different glycoconjugate types in CRISPR edited HEK 293 cells lacking individual or a combination of glycoconjugate types (Narimatsu *et al*., 2019). Here, the authors observed reduced binding for human H1 and H3 HAs in HEK DKO cells, yet no difference in avian H5 HA binding. These findings correlated with the reduced levels of H1N1-GFP but not H5N1-GFP infection observed in our A549 [NO]^-^ and [NG]^-^ DKO cells, indicating that H1N1 viruses showed a preference for N-glycans (Figure 3A). In the same study, the authors observed reduced binding of both human and avian HA subtypes in HEK 293 TKO cells (Narimatsu *et al*., 2019). We also observed reduced binding of H1 and H5 HA proteins in A549 [NOG]^-^ TKO cells as compared to WT cells (Figure 5B); interestingly, our virion binding assays showed significantly lower levels of H1N1 virion binding as compared to H5N1 in A549 [NOG]^-^ TKO cells, suggesting that avian H5N1 virus has the ability to bind to other available glycoconjugate types (20% vs 55%; Figure 5C). Similarly, in single-cycle infection assays, we observed negligible H1N1-GFP infection (<3%) yet robust H5N1-GFP infection (>75%) in A549 [NOG]^-^ TKO cells, indicating that avian H5N1 virus can utilize other types of glycoconjugates as receptors. These differences in glycoconjugate receptor requirements for human versus avian IAV strains were most pronounced in multi-cycle replication assays in [NOG]^-^ TKO cells. Here, we observed a 1-3 log decrease in viral titers for human H1N1 and H3N2 viruses as compared to WT cells, yet little to no change in the replication of several avian IAV strains (Figures 5E and 6A-D). The robust replication of avian IAV strains observed in [NOG]^-^ TKO cells occurred in a Sia-dependent manner, as both pretreatment with sialidase or addition of Oseltamivir treatment decreased H5N1 infection or replication, respectively (Figure 6E-F). Together, these studies demonstrate that avian IAV strains can utilize a broader repertoire of glycoconjugate receptors as compared to human IAV strains for host cell infection.

For truncation of O-glycans, we targeted the biosynthesis gene *C1GALT1* to disrupt the biosynthesis of Core 1 and Core 2 structures, which were the major types of O-glycans detected in our mass spectrometry analysis of WT A549 cells (Figure 4B); however, it should be noted that the expression of sialyl Tn antigen was higher in [NOG]^-^ TKO cells as compared to WT cells (Figure S5B). A recent NMR study demonstrated that avian H5N1 viruses displayed strong interactions with GalNAc moieties in O-glycans, and thus it is possible that avian IAV strains utilize sialyl Tn antigens as entry receptors in [NOG]^-^ TKO cells (Mayr et al., 2018). As previously reported by CFG glycan array studies, it is likely that human IAV strains are unable to efficiently attach to shorter sialoglycans like sialyl Tn antigens (de Vries et al., 2014; Thompson and Paulson, 2021). Together, these findings indicate that extended sialoglycans are dispensable for avian IAV infection, yet may be necessary for efficient infection of human and swine IAV strains. Alternatively, as truncation of multiple glycan types in [NOG]^-^ TKO cells reduced the cell surface availability of Sia, it is possible that human IAV strains require a higher threshold for Sia receptor density as compared to avian IAV strains for host cell binding and infection. This interpretation is in agreement with our previous study in which avian H5N1 virus showed robust replication in A549 cells cultured in the presence of 3Fax-Neu5Ac, a competitive inhibitor of sialyltransferases that decreases the cell surface Sia density (Han et al., 2018). In contrast, H1N1 and H3N2 viruses showed a >3 log reduction in viral titers in 3Fax-Neu5Ac treated A549 cells. Thus, our studies demonstrate that the receptor requirements of avian versus human IAV strains extend beyond the well-established Siaα2,3 vs Siaα2,6 linkage preferences.

In summary, our studies in CRISPR glycoengineered human lung epithelial cells highlights the stark differences in glycoconjugate receptor usage by avian versus human and swine IAV strains. Human IAV strains required one of the three major types of glycoconjugates for efficient host cell infection, with some preference for N-glycans. In contrast, avian IAV strains demonstrated versatility in glycoconjugate receptor usage, as these strains replicated efficiently in cells with truncated N-glycans, O-glycans, and GSL. Taken together, our data indicates that human IAV strains may rely on a limited repertoire of glycoconjugates for host cell infection, whereas avian IAV strains can utilize diverse glycoconjugates for viral entry. These findings will have important implications for our understanding of how glycoconjugate receptor usage determines the host range of zoonotic IAV strains.

## Materials and Methods

### Cells and Viruses

Human lung epithelial (A549) cells and African green monkey kidney (Vero) cells were cultured in DMEM (Gibco) supplemented with 10% fetal bovine serum (FBS, Atlanta Biologicals) and 1% penicillin/streptomycin (P/S). Nuli-1 cells were cultured on Collagen IV coated plates as previously described in Bronchial Epithelial Cell Growth Media (BEGM) with supplements (Lonza) (Zabner et al., 2003). Madin-Darby canine kidney (MDCK) cells were cultured in MEM supplemented with 10% FBS and 1% P/S.

IAV strains used in this study were obtained from different sources – A/Puerto Rico/8/1934 (H1N1, Mount Sinai), high virulent PR8 (hvH1N1 provided by Dr. Georg Kochs), A/Hong Kong/1/1968 (HK68, H3N2), A/Wyoming/03/2003 (H3N2), A/Victoria/3/1975 (H3N2), A/Philippines/2/1982 (H3N2), A/Swine/Minnesota/37866/1999 (SwH1N1), a low pathogenic version of A/Vietnam/1203/2004 (H5N1-low pathogenic), low pathogenic version of A/Netherlands/213/2003 (H7N7-low pathogenic, kindly provided by Dr. Ron Fouchier), A/blue-winged teal/Illinois/10OS1563/2010 (H4N6), A/Rhea/North Carolina/39482/1993 (H7N1), A/shorebird/Delaware Bay/127/2003 (H9N2), and A/Anhui/1/2013 H7N9 2:6 PR8 reassortant virus (H7N9 (PR8)). H1N1-GFP (PR8) and H5N1-GFP viruses were grown as previously described (Kandasamy et al., 2020; Manicassamy et al., 2010). Influenza B strains used in this study - B/Yamagata/16/1988, B/Nevada/03/2011, B/Texas/06/2011. All influenza viruses were either grown in embryonated eggs or MDCK cells. Influenza viruses were aliquoted and stored at −80°C before titering by plaque assay on MDCK cells using 2.4% Avicel RC-581 (a kind gift from FMC BioPolymer, Philadelphia, PA). Viral plaques were quantified 2-3 days post infection by crystal violet staining. Vesicular stomatitis virus expressing GFP (VSV), kindly provided by Dr. Glenn Barber at the University of Miami, FL, was propagated in Vero cells and titers were determined by plaque assay on Vero cells using 1% methylcellulose (Sigma)(Stojdl et al., 2003).

### Generation of CRISPR KO Cells

*MGAT1* and *C1GALT1* KO A549 cells were generated using the lentiCRISPR v2 (#52961, Addgene) single vector system as previously described with Puromycin selection (Sanjana et al., 2014; Shalem et al., 2014). *MGAT1* single KO cells were used to generate *MGAT1/C1GALT1* DKO cells and *MGAT1/UGCG* DKO cells with the pLentiSpBsmBI sgRNA Hygro vector (#62205, Addgene). *MGAT1/C1GALT1* DKO cells were used to generate *MGAT1/C1GALT1/UGCG* TKO cells with the lentCRISPRv2 neo vector (#98292, Addgene). Primers for sgRNA target sites used here are listed in Table S1. On day 2 post lentivirus transduction, A549 cells were subjected to drug selections for ∼14 days at the following concentrations: puromycin −2ug/ml (Invivogen), hyrgromycin - 800ug/ml (Invitrogen), neomycin (G418) - 800ug/ml (Invitrogen). Clonal knockout cells were isolated by seeding ∼100 cells in a 150mm plate and allowing them to grow as individual colonies. Successful KO clones were initially identified by flow cytometry using lectins, and subsequently confirmed by western blot analysis for loss of MGAT1 and/or C1GALT1 expression. In addition, disruption of the intended target sites was confirmed by Sanger sequencing of the region flanking the sgRNA target as previously described (Table S2) (Han *et al*., 2018). Identified insertions and deletion mutations at the sgRNA target sites are listed in Table S2.

### Virus Infections

For assessment of single cycle virus infections, cells were seeded at a density of 3×10^5^ cells per well in a 12-well plate, and infection with GFP viruses was performed in infection media without the addition of TPCK-treated trypsin. At 16 hours post infection, cells were trypsinized and prepared for flow cytometric analysis. Data was acquired on a BD FACSVerse instrument and analyzed using FlowJo software. For assessment of multi-cycle virus replication, WT and knockout A549 cells were seeded in triplicate at a density of 8×10^5^ cells per well in a 6-well plate. On the next day, cell numbers were measured prior to infection. Cells were washed twice with phosphate buffered saline (PBS) and inoculated with virus at the indicated MOI in infection media (DMEM supplemented with 0.2% bovine serum albumin (BSA) and 0.9 µg/ml TPCK-treated trypsin (Sigma). The inoculum was removed after incubation for 1 h at 37°C, and cells were washed twice with PBS before addition of fresh infection media. Supernatants were collected at the indicated times points and stored at −80°C and viral titers were measured by plaque assay. VSV infections were performed as above with infection media not supplemented with TPCK-treated trypsin, and supernatant titers were assessed by plaque assay on Vero cells using 1% methylcellulose (Sigma).

### Sialidase and Oseltamivir Carboxylate Treatment

For sialidase pre-treatment experiments, cells seeded in 12-well plates were pre-treated with 500 mU/mL α2-3/6/8 sialidase from *Clostridium perfringens* (Sigma-Aldrich) for 3 hours at 37°C before infection. For viral neuraminidase inhibitor treatment experiments, Oseltamivir carboxylate (kind gift from Roche) was added at the indicated concentrations to the infection media after 1 hr virus infection.

### Western Blot Analysis

Whole cell extracts were prepared using RIPA buffer containing protease inhibitors (Roche) and western blot analysis was performed with ∼80ug of total protein as previously described (Han *et al*., 2018). Anti-MGAT1 (ab180578 Rabbit monoclonal) and anti-C1GALT1 (ab237734 Rabbit polyclonal) antibodies for western blot analysis were purchased from Abcam and used at a 1:1000 dilution.

### Lectin and Antibody Staining

Fluorescein labeled *Galanthus Nivalis* Lectin (GNL-FITC, #FL-1241 1:250), Cy3 labeled *Sambucus Nigra* Lectin (SNA-Cy3, #CL-1303, 1:500), and fluorescein labeled *Vicia Villosa* Lectin (VVL-FITC, #FL-1231, 1:500) were purchased from Vector Laboratories. FITC-conjugated Cholera Toxin B subunit (CTB-FITC, #C1655, 1:250) was purchased from Sigma-Aldrich. Anti-Sialyl Tn antibody was purchased from ThermoFisher. Cells were incubated with the fluorescently labeled lectins for 30 min on ice in lectin staining buffer (PBS supplemented with 0.2% BSA and 0.1 mM CaCl_2_) and excess unbound lectin was removed by washing in the same buffer. Data was acquired on a BD FACSVerse flow cytometer and analysis was performed using FlowJo Software.

### HA Binding Assay

Cell surface binding of HA proteins was performed with purified recombinant H1 HA and H5 HA (BEI Resources). Briefly, 1×10^6^ cells were incubated with 5 µg of HA on ice for 1 hr, and the unbound protein was removed by washes with staining buffer (PBS supplemented with 0.2% BSA and 2 mM EDTA). Cells were then fixed with 4% paraformaldehyde for 10 min at RT and washed with PBS. Non-specific secondary antibody binding to cells was blocked using blocking buffer (staining buffer supplemented with 5% normal goat serum and 0.1% Tween 20) for 15 min. The amount of bound HA was measured using anti-H1N1 rabbit sera or anti-H5N1 mouse sera for 1 hr, followed by a secondary goat antibody conjugated with Alexa Fluor 647 (Invitrogen) for 30 min. All blocking and staining steps were performed on ice. Cells were analyzed by flow cytometry on a BD FACSVerse flow cytometer; data analysis was performed using FlowJo Software.

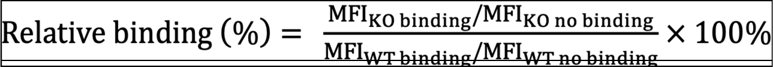

### Cell Surface Virion Binding Assay

Measurements of cell surface binding by IAV particles were performed with 1×10^6^ cells at an MOI=100. Binding of virions was carried out on ice for 1hr 30min, and the unbound virions were removed by extensive washes with staining buffer (PBS supplemented with 0.2% BSA and 2mM EDTA). Cells were then fixed with 4% paraformaldehyde for 10 min at RT and washed with PBS. The levels of virion binding were measured by cell surface staining with anti-H1N1 rabbit sera or anti-H5N1 mouse sera. Briefly, cells were blocked with staining buffer for 15 min, incubated with anti-H1N1 rabbit sera or anti-H5N1 mouse sera (1:500) for 1 hour, washed with blocking buffer to remove unbound antibodies, then subsequently incubated with a secondary goat antibody conjugated with Alexa Fluor 488 (Invitrogen) for 30 min. Data acquisition was performed on a BD FACSVerse flow cytometer and analysis was performed using the FlowJo software.

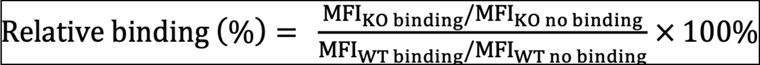

### Glycan Analyses

Analysis for N-glycans and O-glycans as well as quantification of sialic acid was performed at the GlycoAnalytics Core at the University of California at San Diego. For analysis of glycan structures, WT A549 cells and [NOG]^-^ TKO cells were homogenized in the presence of protease inhibitors and proteins were quantified using the BCA kit. Approximately 400ug of proteins were used for the analysis of N- and O-glycans. N-glycans were isolated by treatment with PNGase F (NEB) under denaturing conditions followed by solid phase extraction purification. Approximately 200ug of purified N-glycans were tagged with 2-aminobenzamide fluorophore and analyzed using high pH anion exchange chromatography (HPAEC) analysis. In addition, approximately 140µg of purified N-glycans were subjected to permethylation and analyzed by MALDI-TOF/TOF mass spectrometry. O-glycans were isolated by reductive beta elimination method as previously described and analyzed by MALDI-TOF/TOF mass spectrometry (Bruker Autoflex). Mass spectrometry data was annotated using GlycoWork Bench software.

### Statistical Analysis

Statistical significance was determined by two-tailed unpaired Student’s t test, and pValues ≤ 0.05 were considered significant and denoted with an asterisk. Non-significant values are denoted as ns.

## Acknowledgements

We would like to thank Dr. Adolfo Garcia-Sastre (Icahn School of Medicine) for sharing numerous reagents. Several influenza virus strains and purified HA were obtained from BEI Resources (NIAID). Julianna Han was partly supported by the NIH Molecular and Cellular Biology training program at The University of Chicago (T32GM007183) and the NIH Diversity Supplement (R01AI123359-02S1). Jesse Plung was supported by the National Science Foundation supplement (Award ID 1852070). Miranda Sturtz was supported by the Stinski Undergraduate Research Fellowship (University of Iowa). Dr. Balaji Manicassamy is supported by NIAID grants (R01AI123359 and R01AI127775). The funders had no role in study design, data collection and analysis, decision to publish, or preparation of the manuscript.

## Supplementary Figures

**Figure S1.**
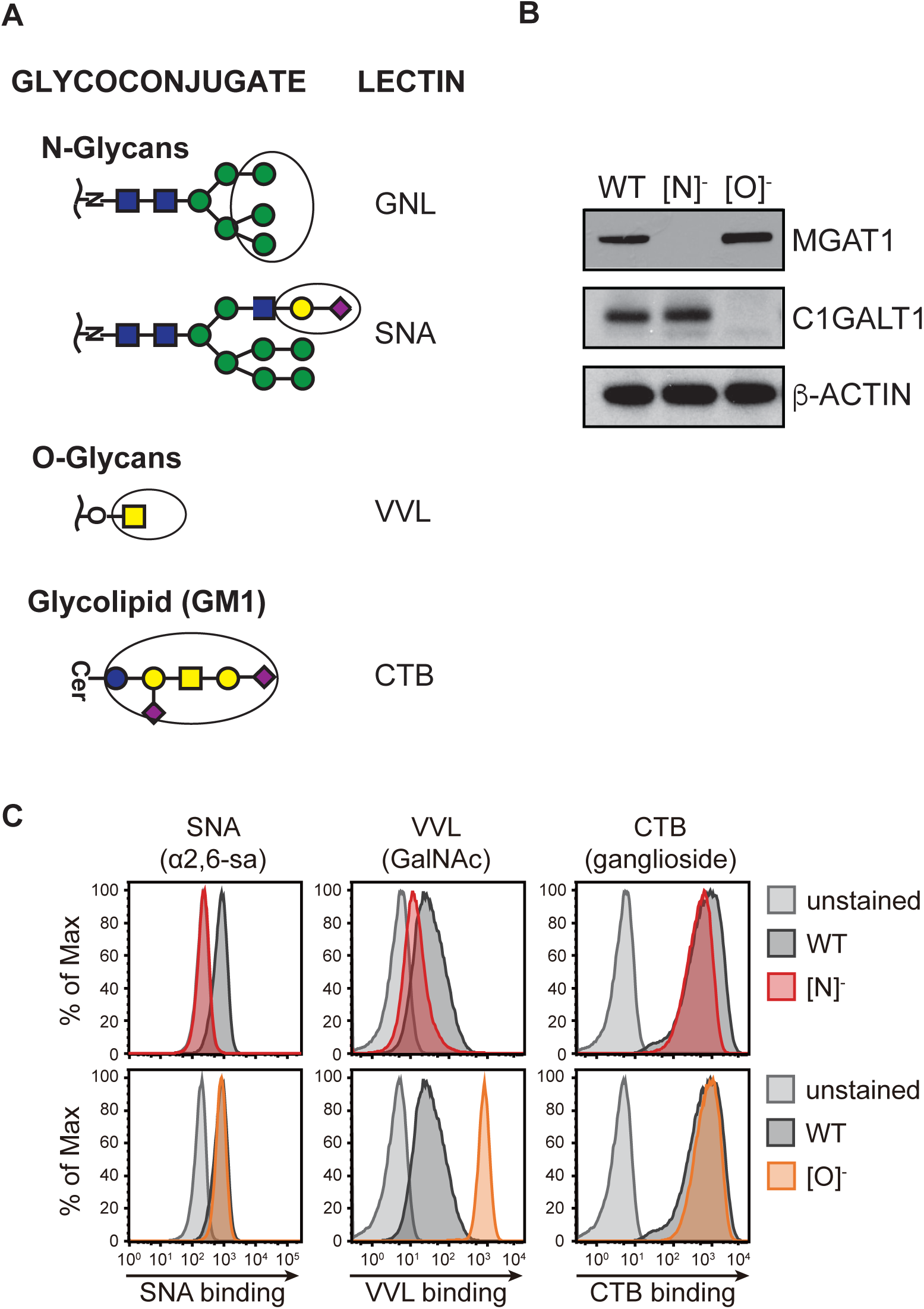
Characterization of [N]^-^ and [O]^-^ KO cells. (A) Binding specificities of different lectins used in this study. GNL - *Galanthus Nivalis* Lectin, SNA - *Sambucus Nigra* Lectin, VVL - *Vicia Villosa* Lectin, and CTB - Cholera Toxin B subunit. (B) Western blot analysis of MGAT1 and C1GALT1 expression in KO cells. β-actin levels are shown as loading controls. (C) Comparison of the binding properties of various lectins in WT and KO cells. Representative flow cytometry plots showing differences in lectin binding. Data are representative of at least two independent experiments.

**Figure S2.**
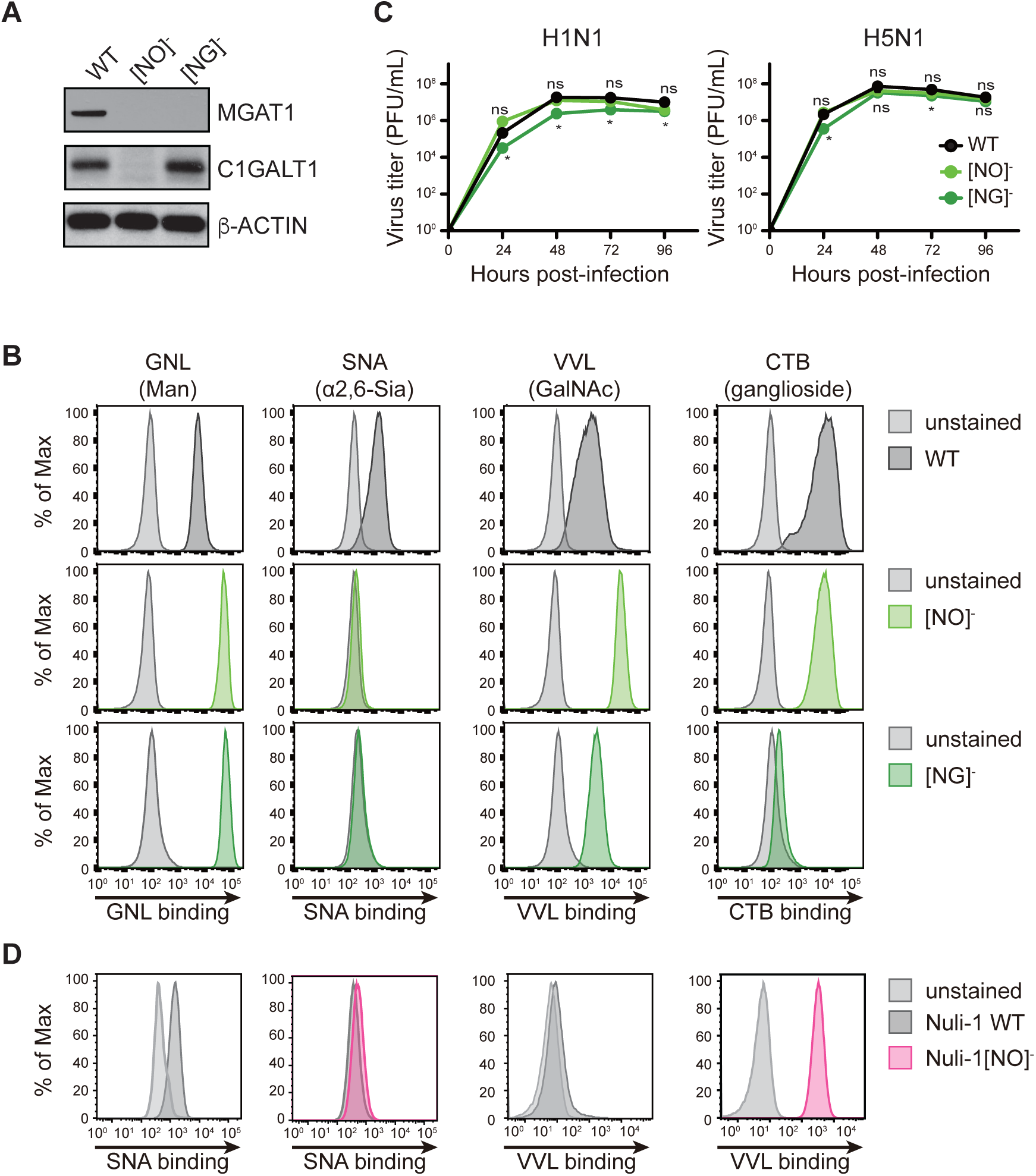
Characterization of [NO]^-^ and [NG]^-^ KO cells. (A) Western blot analysis of MGAT1 and C1GALT1 expression in DKO cells. β-actin levels are shown as loading controls. (B) Comparison of the binding properties of various lectins in WT and DKO cells. Representative flow cytometry plots showing differences in lectin binding. Data are representative of at least two independent experiments. (C) Multi-cycle replication assays with H1N1 and H5N1 in [NO]^-^ and [NG]^-^ DKO cells. WT and DKO cells seeded in 6-well dishes were infected at a low MOI with H1N1 (MOI=0.01) or H5N1 (MOI=0.001) in the presence of TPCK-treated trypsin and viral titers in the supernatants at different hpi were determined by plaque assay in MDCK cells. Data are represented as mean titer of triplicate samples ± SD (PFU/mL). * denotes p-value ≤0.05. ns is non-significant. Data are representative of at least two independent experiments. (D) Comparison of the binding properties of various lectins in Nuli-1 WT and DKO cells. Representative flow cytometry plots showing differences in lectin binding. Data are representative of at least two independent experiments.

**Figure S3.**
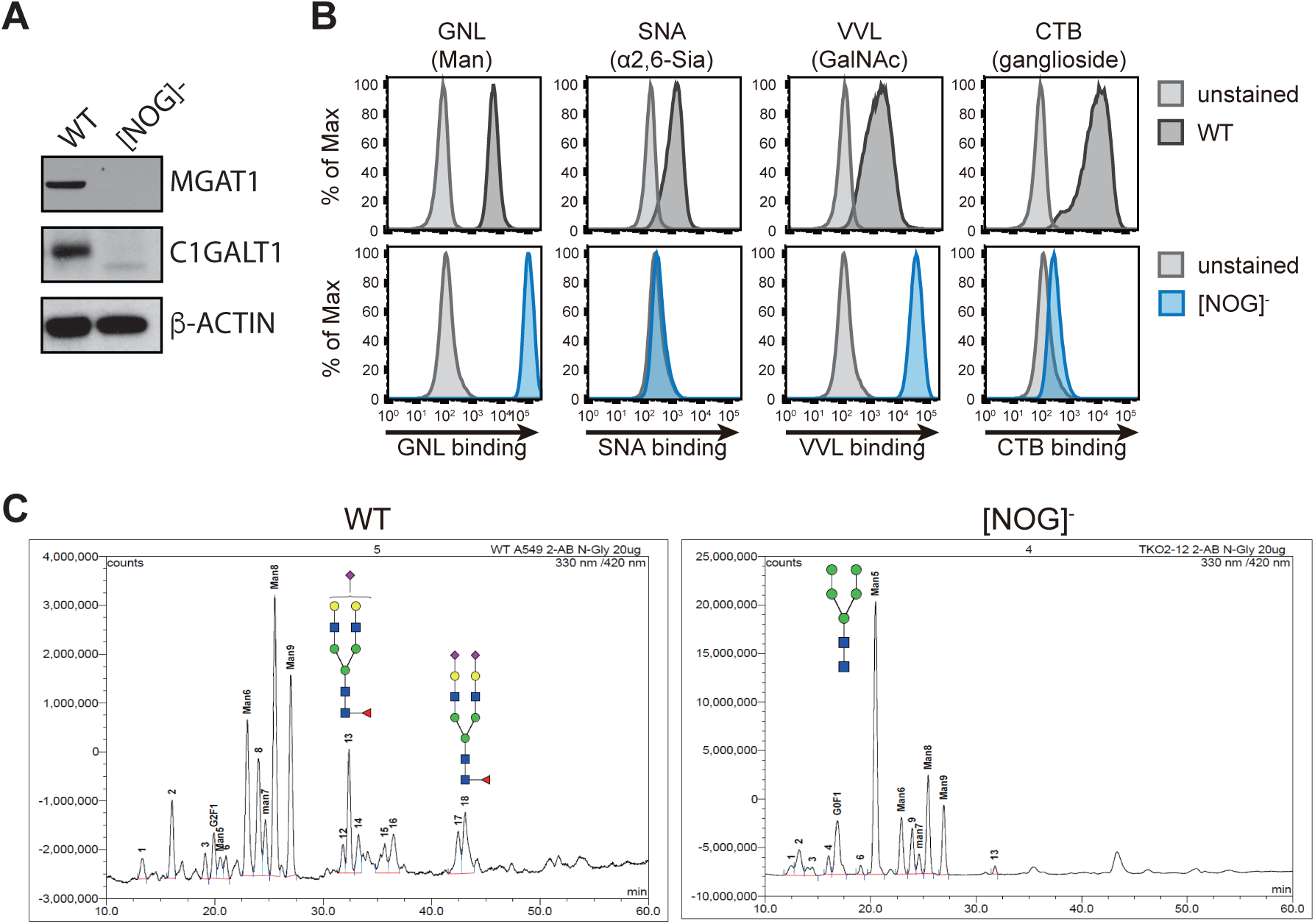
Characterization of [NOG]^-^ TKO cells. (A) Western blot analysis of MGAT1 and C1GALT1 expression in [NOG]^-^ TKO cells. β-actin levels are shown as loading controls. (B) Comparison of the binding properties of various lectins in WT and [NOG]^-^ TKO cells. Representative flow cytometry plots showing differences in lectin binding. Data are representative of at least two independent experiments. (C) Comparison of HPAEC-FL profiles of 2-aminobenzamide (2-AB) tagged N-glycans from WT and [NOG]^-^ TKO cells. N-glycans were isolated by PNGaseF treatment, tagged with 2-AB, and analyzed by HPAEC followed by fluorescence detection.

**Figure S4.**
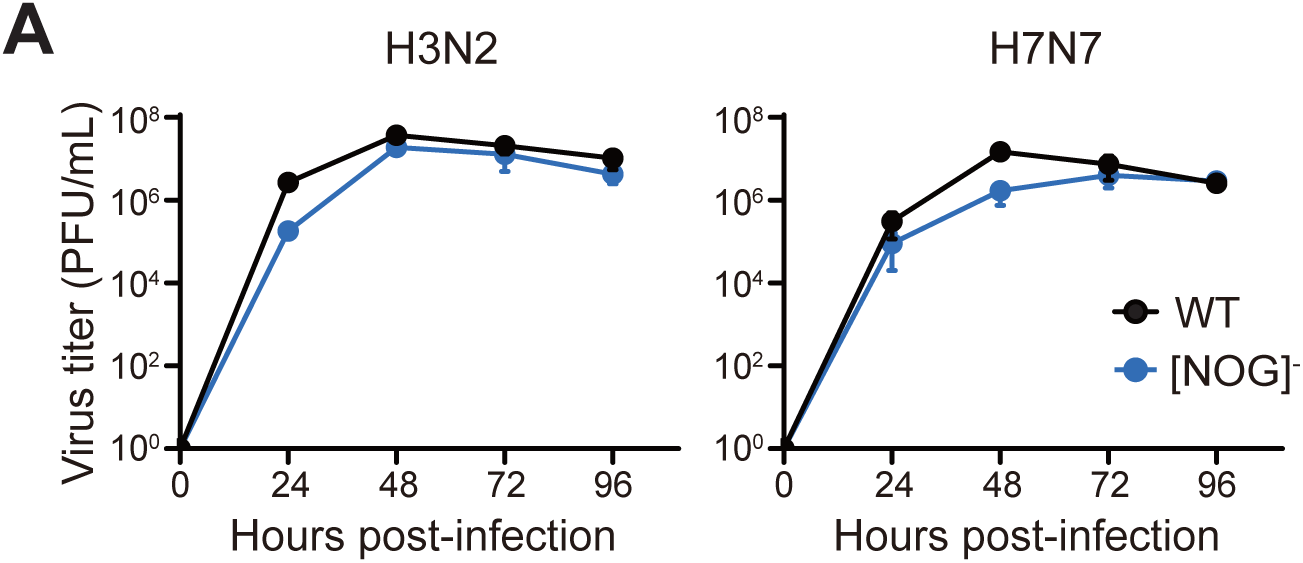
Assessment of H3N2 and H7N7 replication in [NOG]^-^ TKO cells. WT and TKO cells seeded in 6-well dishes were infected at a low MOI with H3N2 or H7N7 (MOI=0.01) in the presence of TPCK-treated trypsin and viral titers in the supernatants at different hpi were determined by plaque assay in MDCK cells. Data are represented as mean titer of triplicate samples ± SD (PFU/mL). * denotes p-value ≤0.05. ns is non-significant. Data are representative of at least two independent experiments.

**Figure S5.**
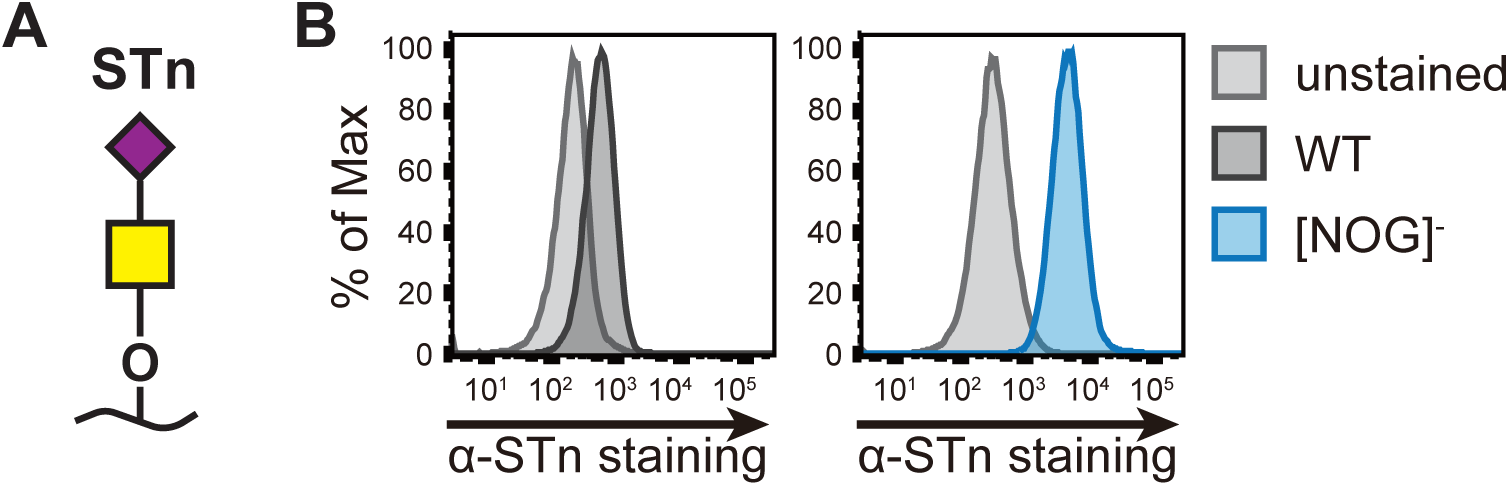
Assessment of Sia levels in [NOG]^-^ TKO cells. (A) Structure of sialyl Tn antigen. (B) Flow cytometry plots comparing STn antigen levels in WT and [NOG]^-^ TKO cells. STn levels were analyzed using anti-STn antibody and representative histograms are shown.

